# Human-exoskeleton interaction force estimation in Indego exoskeleton

**DOI:** 10.1101/2023.03.14.532662

**Authors:** Mohammad Shushtari, Arash Arami

## Abstract

Accurate interaction force estimation can play an important role in optimization human-robot interaction in exoskeleton. In this work, we propose a novel approach for system identification of exoskeleton dynamics in presence of interaction forces as a whole multi-body system regardless of gait phase or any assumption on human-exoskeleton interaction. We hanged the exoskeleton through a linear spring and excited the exoskeleton joints with chirp commands while measuring the exoskeleton-environment interaction force. Several structures of neural networks have been trained to model the exoskeleton passive dynamics and estimate the interaction force. Our testing results indicated that a deep neural network with 250 neurons and 10 time delays can obtain sufficiently accurate estimation of the interaction force, resulting in 1.23 of RMSE on Z-normalized applied torques and 0.89 of adjusted *R*^2^.

## 1. Introduction

Lower-limb exoskeletons have demonstrated promising results in improving mobility and rehabilitation outcomes for individuals with lower-limb motor impairments[1]. These exoskeletons have contributed to the accessibility of early therapeutic intervention for people suffering from spinal cord injuries or stroke boosting their recovery speed[2,3].

The efficacy of these robotic systems depends on their control strategy, particularly, for partially impaired users as the exoskeleton has to encourage exploitation of the user’s residual motor capacity while providing the minimum assistance required for locomotion[4–6]. This requires the exoskeleton to softly switch between the follower and leader roles depending on the user’s performance[7,8]. Otherwise, considerable physical disagreement emerges in the human-exoskeleton interaction causing the user to yield motion to the exoskeleton to maintain comfort at the cost of hindering their motor recovery[5]. Developing an efficient exoskeleton controller, therefore, is not a trivial task as a critical aspect of this endeavor is to optimize the interaction force, which is the force that is exchanged between the user and the exoskeleton during locomotion. Optimizing the interaction force is essential for improving the comfort, stability, and efficiency of the exoskeleton[9].

Human-exoskeleton interaction force is a key information that can reveal human intentions [10]. This makes exoskeleton controllers capable of adjustng their behavior to match each individual’s specific needs during therapy or everyday life. Accurate measurement of the interaction forces is a challenging problem due to the complex nature of the exoskeleton-user interface as the exoskeleton interacts with the user through multiple contact points, including the feet, shanks, thighs, and the torso, which makes it difficult to isolate and measure the forces that are transmitted between human and robot[11]. Consequently, the interaction torque is required to be computed by subtraction of exoskeleton passive dynamics from the actuation torques (see [1,12]). Alternatively, disturbance observers could provide a more robust estimation of interaction torques with the assumption that the human contribution to the gait is an external disturbance ([13–15]).

Many of the previous work on system identification of lower limb exoskeletons has primarily focused on identifying dynamics during the swing phase of walking[16–18]. These studies have often assumed a simplistic two or three-link inverted pendulum model for each leg, while ignoring the coupling between the hip joints. Unfortunately, these simplifications frequently lead to inaccurate interaction torque estimations, particularly during the stance phase of walking. As a result, to achieve precise interaction torque estimation throughout the entire gait cycle, it is crucial to identify the exoskeleton dynamics as a whole multi-body system.

In contrast to previous work, a comprehensive multi-body model considers the complex interactions between the exoskeleton and the user’s body segments, such as the torso. By incorporating these interactions, the model can provide more accurate estimates of the interaction forces and torques that are exchanged between the exoskeleton and the user throughout the entire gait cycle. In this regard, some recent studies have identified exoskeleton dynamics as a whole multi-body system through a data-driven model that involves exciting the exoskeleton joints with rich torque commands while the exoskeleton is fixed to a rigid supporting platform [19]. However, while this approach is an improvement over the previous methods, it fails to capture the exoskeleton-environment interaction, which is the supporting force that fixes the exoskeleton to the platform. As a result, important information about the exoskeleton-environment interaction is ignored, which can limit the applicability of these methods for certain types of exoskeletons. For example, gait training systems such as Locomat (Hocoma, Switzerland), where the hip torque is applied between the femur and a fixed support [20], are better suited for this method than exoskeletons with trunk segments such as Indego (Parker Hannifin, USA), where the hip torque is applied between the femur and trunk, and the interaction occurs primarily between the user’s core and the exoskeleton trunk segment [21]. Therefore, to accurately identify the exoskeleton’s dynamics and interaction torques, it is necessary to include the exoskeleton-environment interaction in the system identification process.

We propose a method for the identification of the exoskeleton’s passive dynamics that measures the exoskeleton interaction with the environment and includes it in the dynamic identification of the exoskeleton. To make this idea feasible, we limited the exoskeleton-environment interaction through an accurate linear spring allowing us to measure the external forces being applied to the exoskeleton from the spring length while exoskeleton joints are being extensively excited. Using the accurate estimation of the interaction forces, we then identify the dynamics of the exoskeleton with an artificial neural network (ANN).

## 2. Materials and Methods

### 2.1. Interaction Torque Modelling

Figure 1.A illustrates the Indego exoskeleton and its coordination systems in the sagittal plane. This lower limb exoskeleton has 7 degrees of freedom (DOF) including one rotational and two translational DOFs of the trunk (3 DOFs in total) with respect to the global frame and only 4 active DOFs of the exoskeleton knee and hip joints of the right and left legs. The exoskeleton has also two carbon-fiber ankle foot orthoses which were removed from the exoskeleton due to their negligible mass. Given that the Indego exoskeleton does not have joints with off-sagittal-plane degrees of freedom (such as hip abduction and adduction DOFs) the off-sagittal dynamics of the exoskeleton during straight walking is negligible compared to its dynamics in sagittal plane. The Indego lower-limb exoskeleton’s dynamics, with actuated hips and knees, in presence of interaction forces and torques in the sagittal plane is considered as

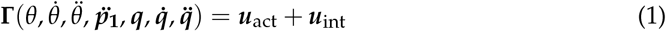

where 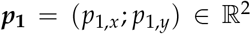 and 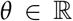 are the exoskeleton’s trunk segment position and orientation, respectively, described in the fixed global frame in the sagittal plane and 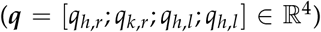 consists of the exoskeleton right hip and knee joint angles followed by the left hip and knee joint angles. ***u***_act_ = [**0**_3×1_; ***τ***_act_] is the applied motor torques at the hip and knee joints of both legs 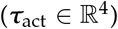. ***u***_int_ = [***f***_int_; *m*_int_; ***τ***_int_] includes the interaction force 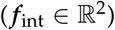 and moment 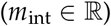 applied to the exoskeleton trunk as well as the interaction torques applied to the exoskeleton joints 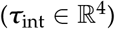. Finally, **Γ**(.) represents the exoskeleton passive dynamics containing the inertial forces, centrifugal and Coriolis forces, and the forces due to gravity [22]. We note that ***p*_1_** and ***ṗ*_1_** do not appear in **Γ** as there are no forces being applied to the trunk due to the trunk linear position (***p*_1_**) and velocity(***ṗ*_1_**). Figure 1.A provides a visual representation of the Indego lower-limb exoskeleton.

**Figure 1.**
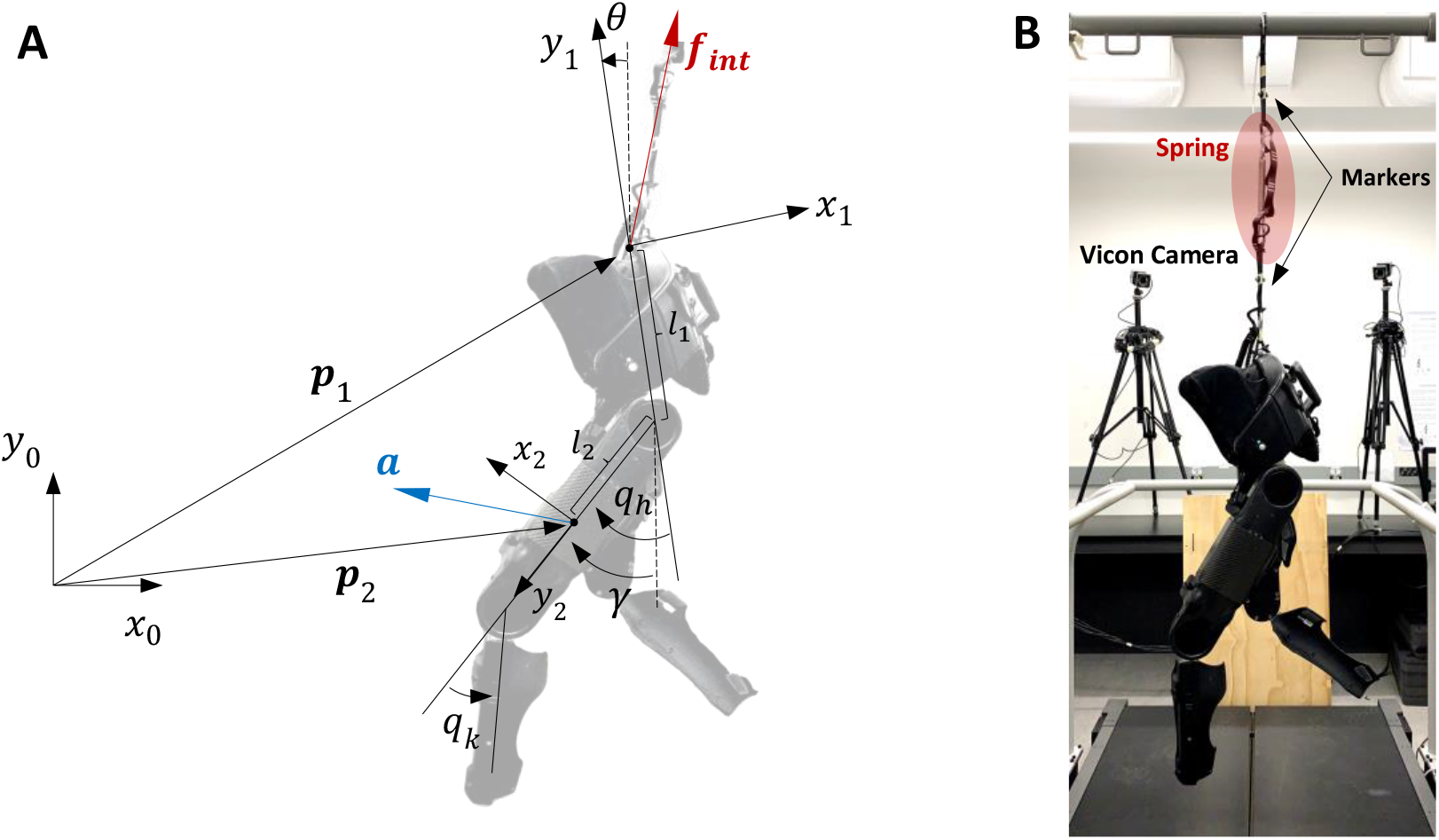
(**A**) The exoskeleton coordinates and joint angles. The global, trunk and thigh frames are denoted by 0, 1, and 2 indices, respectively. Trunk and thigh angles with respect to gravity vector are denoted by *θ* and *γ*, respectively. *q_h_* and *q_k_* represent the hip and knee joint angles, respectively. ***p*_1_** and ***p*_2_** shows the position of the trunk and thigh coordinated in the global frame ((*x*_0_, *y*_0_)). In blue, the acceleration vector (***a***) measured by the accelerometer embedded in the exoskeleton’s thigh is plotted against the thigh frame. In red, we plot the interaction force vector (***f_int_***) calculated from the linear spring’s deflection measured by motion capture. The distance from the trunk frame origin to the hip joint rotation axis and from there to the accelerometer location on the exoskeleton’s thigh is indicated by *l*_1_ and *l*_2_ and gravity axis is illustrated by dashed lines. (**B**) The exoskeleton in hung position during data collection for dynamics identification. Joints are excited during the experiment while the exoskeleton interaction with the environment is limited to the force applied through a linear spring with reflective markers on both ends enabling us to measure spring deflection using a motion capture system and, consequently, the applied force.

### 2.2. System Identification

To estimate the interaction forces and torques (**u**_int_), we first identify the exoskeleton passive dynamics (**Γ**) by training an ANN on a dataset collected by exciting the exoskeleton joints in a condition at which we are able to measure the exoskeleton interaction forces with the environment. This was accomplished by hanging the exoskeleton from the ceiling by a spring with K = 1926.4 N/mm with reflective markers attached to both ends (Figure 1.B). This way, we could measure the spring deflection and consequently, compute the ***f***_int_ accurately using a motion capture system consists of eight Vero Cameras (Vicon, UK).

Exoskeleton joints were excited in three different scenarios each for 500 seconds by applying chirp command trajectories (controlled by joint-level PD controllers with *k_p_* = 1.5 N.m/deg and *k_d_* = 0.1 N.m.s/deg) with unique initial and final frequencies to prevent phase lock between joints (see Figure 2). The chirp signals were designed to cover at least 80% of each joint range of motion with frequencies between 0.2 to 0.9 Hz covering the speed ranges bellow 1.4 m/s for healthy individuals [23]. The Spring force measurement as well as the exoskeleton data consisted of its joint angles (***q***), angular velocity 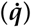, the angle of each thigh with respect to gravity 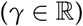, linear acceleration of each thigh 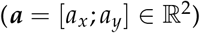, and the applied joint torques (***τ***_act_) were acquired at 200Hz. By choosing different up-chirp and down-chirp rates for the command trajectories to each joint, we applied three different patterns for interaction forces to the exoskeleton trunk. During the first scenario (Train dataset), the right hip and left knee frequencies up-chirped from 0.2 Hz to 0.9 Hz, while the left hip and right knee frequencies down-chirped from 0.9 Hz to 0.2 Hz. All joints in the second scenario (Validation) up-chirped from 0.2 Hz to 0.9 Hz. Finally, in the last scenario (Test), the joints on the right leg up-chirped from 0.2 to 0.9 Hz while those on the left leg down-chirped from 0.9 to 0.2 Hz. In all the cases the frequency sweep rate for the up-chirping or down-chirping joints were selected slightly different to prevent phase locking between joints (Figure 2).

**Figure 2.**
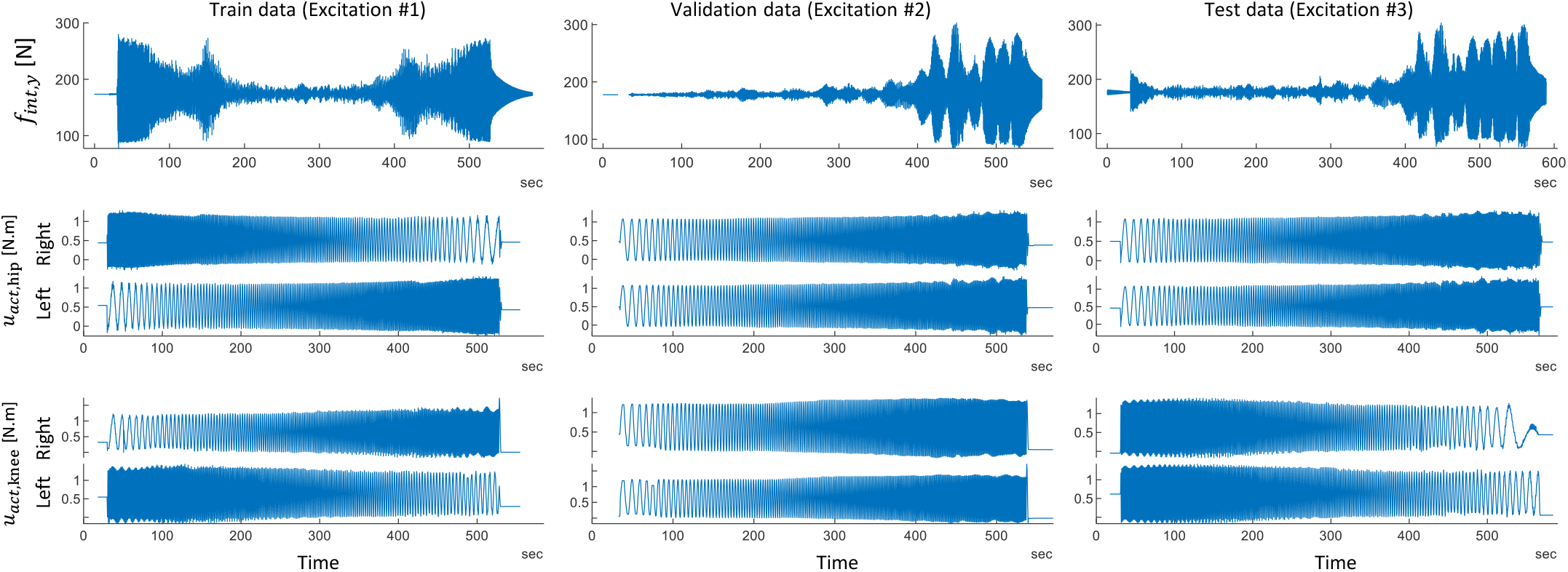
The hip and knee actuation torques applied to the left and right exoskeleton legs, as well as the interaction force in the y axis. During the generation of each of the training, validation, and test datasets, exoskeleton joints were activated with unique and different chirp commands resulting in different exoskeleton-environment interaction profiles.

According to Equation 1, we then compute the trunk orientation (*θ*) and its derivative in addition to its linear acceleration 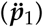 to train a data-driven model for estimation of exoskeleton passive dynamics (Γ). Even though this information is not directly included in the exoskeleton measurements, they are obtained using either of the thigh’s linear accelerometer measurements (***a***) and through obtaining its angle (*γ*) with respect to the gravity vector (*g*). Trunk orientation is then computed using the angle (*γ*) and the corresponding hip joint angle (the angle between the thigh and the upper segment) measured using the motor encoders. Therefore we will drop the leg index hereafter. According to Figure 1.A, we can simply obtain trunk orientation as

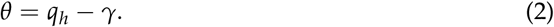

To obtain 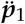, we first mapped thigh acceleration (***a***) from the thigh coordinate to the global coordinate frame. As the thigh coordinate frame ((*x*_2_, *y*_2_)) has rotated clockwise by *π* – *γ* rad compared to the global frame ((*x*_0_, *y*_0_)), the linear acceleration measured in the thigh frame (***a***) can be mapped to the global frame as

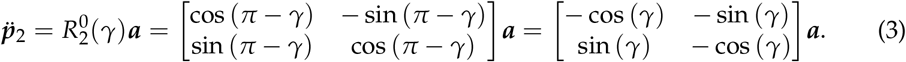

Moreover, The following equation holds between the origin of the trunk frame and the accelerometer position with respect to the global frame (***p***_2_ = (*P*_2,*x*_; *p*_2,*y*_))

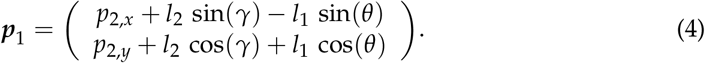

Taking the second derivative from the above equation and substituting Equation 3, we obtain

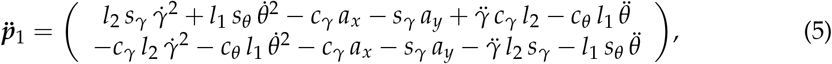

where *c_x_* and *s_x_* are shorthanded versions of *sin*(*x*) and *cos*(*x*), respectively. According to Equation 2 and Equation 5, both *θ* and its derivatives, as well as 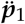, could be obtained as a function of exoskeleton joint angles (***q***), thigh accelerations (***a***), thigh orientation with respect to gravity (*γ*), and their derivatives. In other words, we have

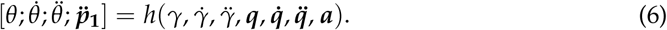

Henthforce, without loss of generality, we can consider Γ only a function of exoskeleton measurements and rewrite Equation 1 as

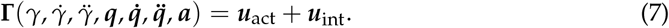

Moreover, we define the origin of the exoskeleton’s trunk frame exactly at the point where the spring force is applied (Figure 1.A). This allows us to consider *m*_int_ = 0. Equation 1 can be, thus, further simplified as

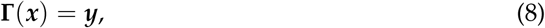

where 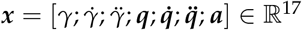 and 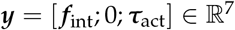.

Various Time-Delay ANNs [24] with tanh activation function were trained on the training dataset to learn the exoskeleton’s passive dynamics 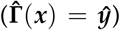 by minimizing the root mean squared error (RMSE) between the Z-score normalized measured (***y***) and estimated (***ŷ***) forces and torques using the scaled conjugate gradient backpropagation algorithm [25]. The training was stopped when 10 consecutive iterations has not improved the performance of the ANN on the validation dataset. Different combinations of network structure (ranging from a single hidden layer with 10 neurons to three hidden layers with 250 neurons in total), number of input delays (ranging from 0 to 50 samples), and *L*^2^ weight regularization factors (ranging from 0 to 0.5) were evaluated according to (Figure 3.A), by training each network five times and computing their mean performance on the validation dataset to compare networks and to obtain the best architecture.

**Figure 3.**
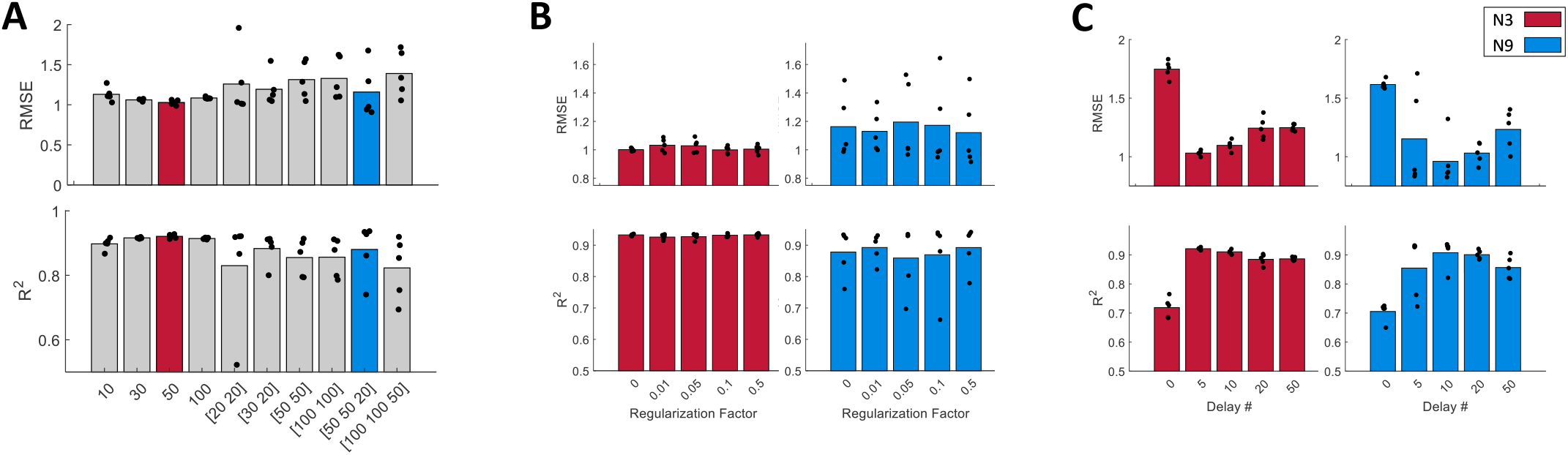
(**A**) Validation performance of ANNs with 10 different structures in terms of RMSE and adjusted *R*^2^ computed on the z-normalized data. The training procedure was repeated 5 times for each structure to study the sensitivity of the training procedure to the network’s initial weight for each structure. Each validation performance is denoted by black dot while the average performance is denoted by bars. Training was done with 5 input delays and no regularization. N3 (denoted by red) and N9 (denoted by blue) demonstrate better performance compared to the other networks. N9, however, exhibits more variability across training repetitions. Those networks are selected for further analysis to investigate the effect of regularization factor and the number of input delays. (**B**) Comparison of N3 and N9 performance across different selection of regularization factor. The variation of the N9 performance across training repetitions is unaffected by regularization. (**C**) Effect of the number of input delay on the N3 and N9 networks performances. N3 has the best performance with 5 input delays while N9 reaches the best performance by at 10 input delays.

## 3. Results and Discussion

### 3.1. Tuning the network structure and hyperparameters

To tune the network structure and complexity, we trained ten networks with complexities ranging from a single hidden layer with 10 neurons up to a network with 3 hidden layers with 250 neurons in total. Figure 3.A shows the average RMSE and the coefficient of determination (*R*^2^) computed using each of the trained networks on the validation datasets. The training procedure is repeated 5 times for each network to account for the randomness of the training process without regularization coefficient and with input delays equal to 5. Our results indicated that the average test performance drops as the network complexity increases. This is indicated by the increase in the average RMSE and the decrease in the average adjusted *R*^2^ after N3 (the third network denoted by red in Figure 3.A) which resulted in one of the best performance in consistently in all 5 training trials (RMSE = 1.03 *R*^2^ = 0.92). On the other hand, the RMSE and *R*^2^ variations on the validation dataset also increase with the network complexity. This can be explained by the association between the networks expressivity and overfitting. N9 (ninth network denoted by blue in the same graph), for example, obtained one of the best validation performances among all networks 3 out of 5 times. To further improve the results, the training procedure was repeated for N3 and N9 with regularization coefficients ranging from 0 to 0.5 and with different numbers of input delays ranging from 0 to 50 samples equal to 0-250 ms.

Figure 3.B investigates the effect of the regularization coefficient on N3 and N9 performances indicating that increasing the regularization coefficient improves neither the network performance in the case of N3 nor the consistency of the validation results in case of N9. The number of input delays, on the contrary, shows to be effective in enhancing both network performances. According to Equation 8, providing ANNs with histories of ***x*** will not be effective as **Γ** is solely a function of ***x***. However, the best performance for N3 is obtained with 5 input delays (RMSE = 0.99 and *R*^2^=0.92) while N9’s best performance is obtained with 10 input delays (RMSE = 0.82 and *R*^2^ = 0.93). The underlying reason is that by adding input delays, the ANNs can learn to smooth the input data and therefore become less sensitive to the input noise. The more flexible structure of N9 enables it to benefit from the additional information embedded in the input history while N3 capacity becomes saturated with more than 5 input delays, and therefore, N9 is able to reach a less biased performance. Hence we chose N9 consisting of three hidden layers each with 50, 50, and 20 neurons, respectively with 10 input delays for estimation of the exoskeleton dynamics.

### 3.2. N9 performance on test dataset

The overall RMSE and *R*^2^ of N9 on the test dataset are 1.23 with *R*^2^ = 0.89, respectively. Figure 4 shows the test performance of N9 for each output channel. The relative error is computed by normalizing the estimation error over the maximum range of each output channel. For example, according to Figure 2 the range of *f_int,y_* is about 200 N. In this case, 1% of relative error means 2N of force estimation error. The trained network exhibits an accurate estimation of applied exoskeleton joint torques with a coefficient of determination greater than 0.9. The knee torque estimation is slightly more biased compared to the hip due to the more pronounced static friction at the knee. Particularly, at lower speeds, the larger mass (4.7 Kg) of the thigh segment dominates the static friction of the motor gears at the hip joint resulting in more predictable dynamical behaviour. The light weight of the shank segments, in contrast, is negligible (0.6 Kg) compared to the static friction and motor backlash causing a greater estimation bias at the knee. This is evident from the dead zone-like behaviour emerged in the *y*-*ŷ* graphs for the knees. At the higher speeds when static friction is dominated by inertial and damping torques, the knee torque estimations are more accurate than the hip joint due to its smaller moment of inertia leading to a higher coefficient of determination at the knee joint. The RMSE value for the interaction forces is higher than the joint torques. Simplifying the exoskeleton dynamics into the sagittal plane is the main reason behind this as our model cannot represent the applied interaction force to the exoskeleton trunk in the *z* direction. The external forces are also estimated with an RMSE smaller than 1.6 with negligible bias (ME<0.2%).

**Figure 4.**
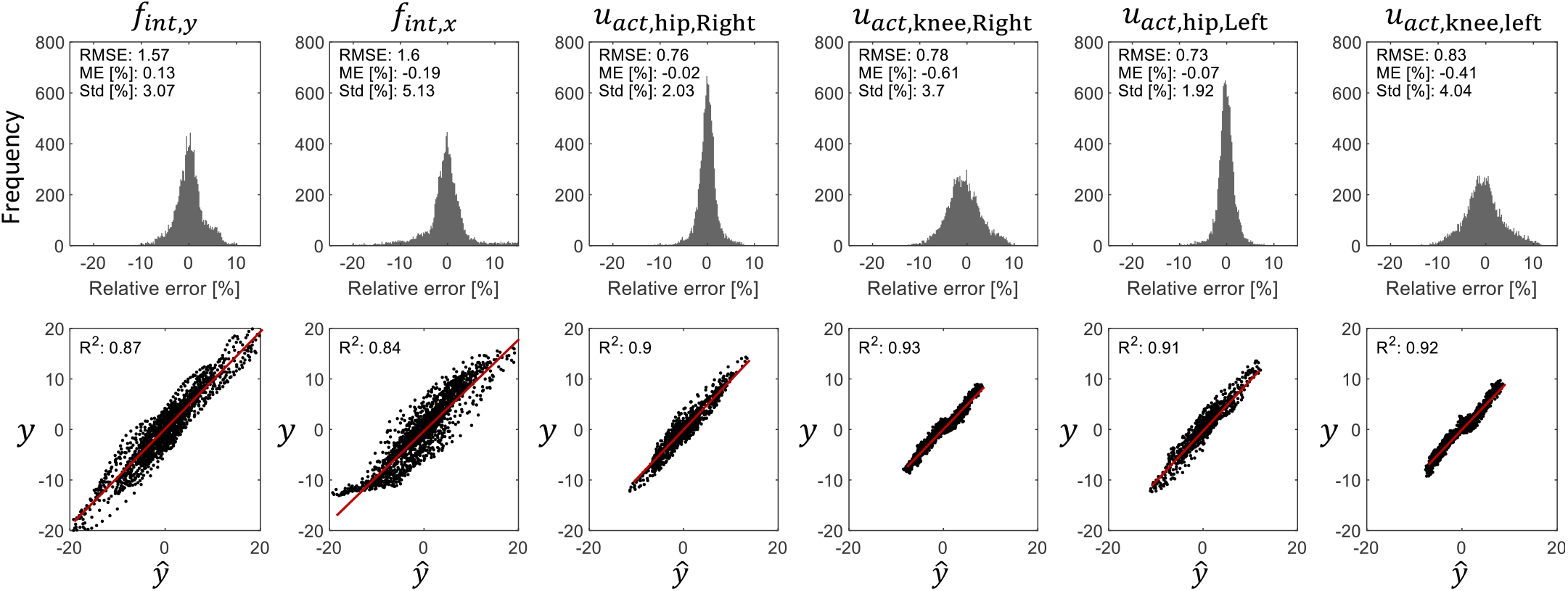
The Test performance of N9 for each output channel. In the top row, distribution of the estimation error of external force or joint applied torques relative to their peak-to-peak range is depicted. Interaction forces in the *y* and *x* direction exhibit higher estimation error compared to those of the exoskeleton active joints. Applied torques to the hip joints, in particular, is estimated with higher accuracy smaller standard deviation. Even though knee joint applied torques are estimated with similarly small errors (small bias), they show higher standard deviation associated to the more prominent role of static fruition. The bottom row shows the correlation between the measured data and the N9 estimation for each output channel. The estimated outputs have relatively lower coefficient of determination in case of interaction forces while they show *R*^2^>0.9 for the active joints of the exoskeleton.

## 4. Conclusion

The dynamics of a lower-body exoskeleton is identified and used for the estimation of the human-exoskeleton interaction torques. Exoskeleton is hung using a linear spring allowing us to measure the exoskeleton-environment interaction forces by motion capture. Meanwhile, exoskeleton joints were excited with different chirp commands designed to cover the range of motion of all joints in three scenarios, each used either for, train, validation, or test of the data-driven model. Various time-delay ANNs were trained and validated on the collected datasets and the network structure, the number of input delays, and regularization factors were tuned according to their validation performances. Testing results indicate accurate performance in exoskeleton joint torque estimation (RMSE <0.85, ME<0.6%, and *R*^2^>0.9) as well as acceptable performance on interaction force estimation (RMSE<1.6, *R*^2^>0.84). Modelling the exoskeleton motion in the sagittal plane is one of the limitations of this work. Even though a high accuracy in joint torque estimation is obtained, with more accurate modelling of the exoskeleton in 3D space, we can model the interaction forces applied to the exoskeleton better. This requires the development of a drift-less orientation estimator enabling us to monitor the orientation of the exoskeleton trunk in 3D space continuously. This direction will have implications for building a data-driven ground reaction force estimator in our future works. Our next step, however, is to integrate the identified exoskeleton dynamics in our adaptive trajectory and feed-forward torques controllers [26,27] which are tested only in simulation due to the lack of ability in the estimation of human-exoskeleton interaction torques.

## Disclaimer/Publisher’s Note

The statements, opinions and data contained in all publications are solely those of the individual author(s) and contributor(s) and not of MDPI and/or the editor(s). MDPI and/or the editor(s) disclaim responsibility for any injury to people or property resulting from any ideas, methods, instructions or products referred to in the content.

